# Slippery when wet: cross-species transmission of divergent coronaviruses in bony and jawless fish and the evolutionary history of the *Coronaviridae*

**DOI:** 10.1101/2021.03.22.436364

**Authors:** Allison K. Miller, Jonathon C.O. Mifsud, Vincenzo A. Costa, Rebecca M. Grimwood, Jane Kitson, Cindy Baker, Cara L. Brosnahan, Anjali Pande, Edward C. Holmes, Neil J. Gemmell, Jemma L. Geoghegan

**Affiliations:** Department of Anatomy, University of Otago, Dunedin, New Zealand; Department of Biological Sciences, Macquarie University, Sydney, Australia; Department of Microbiology and Immunology, University of Otago, Dunedin, New Zealand; Kitson Consulting Ltd, Invercargill/Waihopai, New Zealand; National Institute of Water and Atmospheric Research, New Zealand; Animal Health Laboratory and Diagnostic and Surveillance Directorate, Ministry for Primary Industries, Upper Hutt, New Zealand; Marie Bashir Institute for Infectious Diseases and Biosecurity, School of Life and Environmental Sciences and School of Medical Sciences, The University of Sydney, Sydney, NSW 2006, Australia; Institute of Environmental Science and Research, Wellington 5018, New Zealand

**Keywords:** meta-transcriptomics, virus discovery, fish, evolution, phylogeny, Coronaviridae, lamprey, coronavirus

## Abstract

The *Nidovirales* comprise a genetically diverse group of positive-sense single-stranded RNA virus families that infect a range of invertebrate and vertebrate hosts. Recent metagenomic studies have identified nido-like virus sequences, particularly those related to the *Coronaviridae*, in a range of aquatic hosts including fish, amphibians and reptiles. We sought to identify additional members of the *Coronaviridae* in both bony and jawless fish through a combination of total RNA sequencing (meta-transcriptomics) and data mining of published RNA sequencing data, and from this reveal more of the long-term patterns and processes of coronavirus evolution. Accordingly, we identified a number of divergent viruses that fell within the *Letovirinae* subfamily of the *Coronaviridae*, including those in a jawless fish – the pouched lamprey. By mining fish transcriptome data we identified additional virus transcripts matching these viruses in bony fish from both marine and freshwater environments. These new viruses retained sequence conservation in the RNA-dependant RNA polymerase across the *Coronaviridae*, but formed a distinct and diverse phylogenetic group. Although there are broad-scale topological similarities between the phylogenies of the major groups of coronaviruses and their vertebrate hosts, the evolutionary relationships of viruses within the *Letovirinae* does not mirror that of their hosts. For example, the coronavirus found in the pouched lamprey fell within the phylogenetic diversity of bony fish letoviruses, indicative of past host switching events. Hence, despite possessing a phylogenetic history that likely spans the entire history of the vertebrates, coronavirus evolution has been characterised by relatively frequent cross-species transmission, particularly in hosts that reside in aquatic habitats.

## Introduction

The *Coronaviridae* are a family of enveloped, positive-strand RNA viruses that have a broad host range, although they are predominantly associated with mammals and birds. Several coronaviruses have emerged as highly pathogenic viruses in humans and other animals, the most recent and notable of which is *Severe acute respiratory syndrome coronavirus-2* (*SARS-CoV-2*) (Wu et al. 2020) which, at the time of writing, has infected over 100 million people since its initial identification at the end of 2019. Coronaviruses, like other families within the order *Nidovirales*, possess the longest genomes seen in RNA viruses (27-34 kb in the case of the coronaviruses), and seemingly replicate with higher fidelity than other known positive-sense RNA viruses due to RNA proofreading associated with an exonuclease domain (Denison et al. 2011).

Most coronaviruses are classified within the sub-family *Orthocoronavirinae* that infect mammals and birds, as well as a single virus identified in a reptilian host (Shi et al. 2018). However, our understanding of the host range of the *Coronaviridae* has recently been enhanced by large-scale metagenomic studies, including the identification of a distinct second sub-family - the *Letovirinae*. Specifically, an analysis of published transcriptome data identified a novel coronavirus in the ornamented pygmy frog (*Microhyla fissipes*) termed *Microhyla letovirus* (Bukhari et al. 2018). A subsequent virological survey of dead and moribund cultured Chinook salmon (*Oncorhynchus tshawytscha*) identified *Pacific salmon nidovirus* (Mordecai et al. 2019). Phylogenetic analysis revealed that *Microhyla letovirus* and *Pacific salmon nidovirus* formed a sister group to all other known coronaviruses, with the large genetic divergence between them suggesting that this group is likely far larger and under-sampled (Bukhari et al. 2018; Mordecai et al. 2019). It is therefore unsurprising that more recent probes of public sequencing data have identified additional coronavirus-like sequences in non-mammalian aquatic vertebrate hosts, namely in fish, reptile and amphibian transcriptomes (Edgar et al. 2020). Indeed, this extensive data mining study identified six novel viruses within the *Letovirinae*: four in bony fish, one in the axolotl (*Ambystoma mexicanum*) and a further, more divergent virus in the loggerhead sea turtle (*Caretta caretta*) (Edgar et al. 2020).

The central aim of the current study was to determine whether coronaviruses might be present in a wider range of fish hosts, and from this reveal more of their evolutionary history, particularly whether coronaviruses have co-diverged with their hosts over many millions of years or whether there has been frequent cross-species transmission (i.e. host jumping) as underpins the emergence of *SARS-CoV-2*, and whether such cross-species transmission occurs in some hosts more frequently than others. To this end, we screened for coronaviruses using total RNA sequencing (i.e. meta-transcriptomic) data from bony and jawless fish sampled in New Zealand and Australia, combined with data mining of published RNA sequencing data sampled from fish within the National Centre for Biotechnology Information (NCBI) Sequence Read Archive (SRA). This enabled us to identify new coronavirus sequences that not only significantly expanded the phylogenetic diversity of the coronaviruses, but that provided important new insights into the fundamental patterns and processes of viral evolution in this virus family.

## Results

### Identification of a novel coronavirus in the common carp

As part of a large virological survey of freshwater fish across the Murray-Darling Basin in Australia, ten species of native and invasive bony fish were caught and subjected to meta-transcriptomic analysis (Costa et al. 2021). This led to the identification of a novel member of the *Coronaviridae*, termed *Murray-Darling carp letovirus* (Costa et al. 2021). This partial virus sequence was obtained from common carp sampled from Edward River, New South Wales. This divergent viral sequence shared ORF1b sequence similarity with *Pacific salmon nidovirus* (51%), as well as with various gammacoronaviruses, including *Bottlenose dolphin coronavirus* and *Beluga whale coronavirus* (46%), and had a standardised abundance of 1.2 × 10^−8^ (Table 1). The phylogenetic position of this virus is discussed in more detail below and elsewhere (Costa et al. 2021).

**Table 1.**
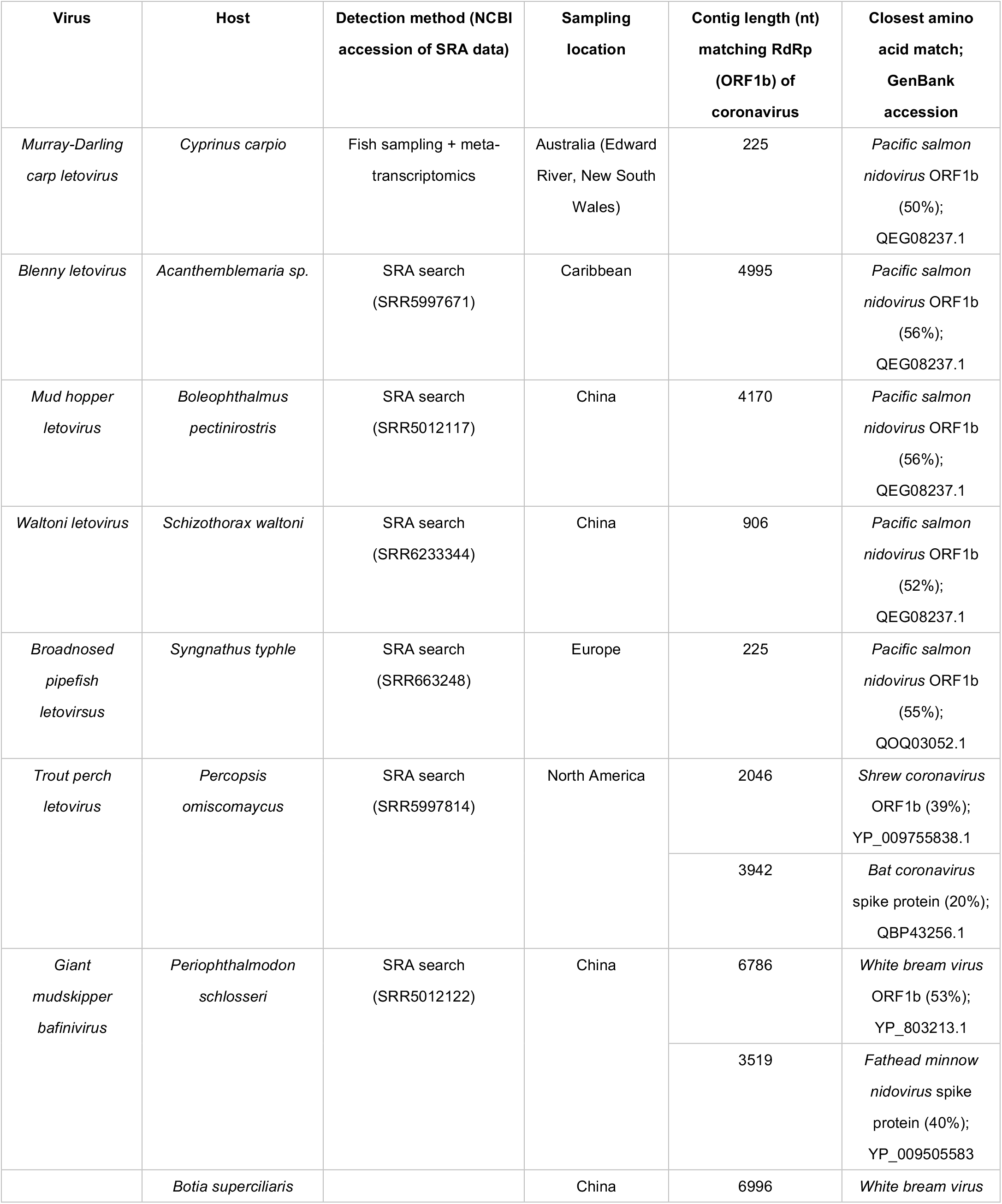
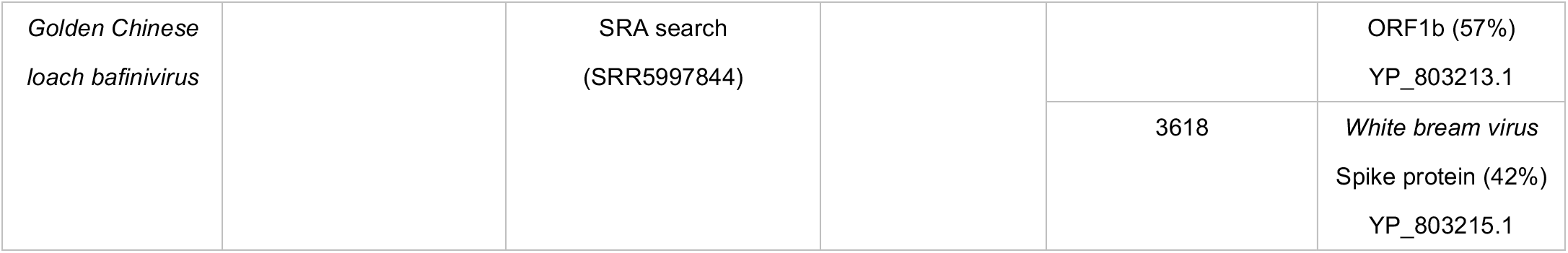
Novel viruses identified in this study from bony fish in Australia and by mining the Sequence Read Archive (SRA), their detection method and sampling locations.

### Identification of a novel coronavirus in the pouched lamprey

Following a disease investigation into Lamprey Reddening Syndrome (LRS) in pouched lamprey from New Zealand (*Geotria australis*) – te reo Māori: kanakana (South Island), or piharau (North Island) – by the New Zealand Ministry for Primary Industries (MPI) in 2012, 12 individuals (eight from this investigation and four more recently collected control samples) were subjected to meta-transcriptomic analysis. This led to the identification of a novel and divergent member of the *Coronaviridae* in several individuals. We tentatively named this new virus *Kanakana letovirus*, and as each virus shared >97% sequence similarity at the nucleotide level they are likely genomic variants of the same virus species.

We detected sequence reads that shared sequence identity to coronaviruses in seven of the eight LRS-affected individuals (confirmed by RT-PCR). No coronaviruses or other viruses were found in the remaining LRS sample nor the four control samples. However, we were only able to generate consensus virus sequences from three of these due to the low and fragmented virus abundance in the other samples. All three samples possessed virus transcripts with sequence similarity to the open reading frames 1a (ORF1a) and 1b (ORF1b) common to members of the *Coronaviridae*, the latter of which contains the viral RNA-dependent RNA polymerase (RdRp) (Table 2). We also identified transcripts with marked sequence similarity to the *Nidovirales* spike (S) protein. However, the closest sequence match for the S protein transcripts were viruses from *Porcine torovirus* and *Atlantic salmon bafinivirus*, both non-coronavirus members of the *Nidovirales* (family *Tobaniviridae*), albeit these were highly divergent sequences with only ~25-27% amino acid sequence similarity (Table 2). Phylogenetic comparisons of the ORF1ab and S genomic regions of *Kanakana letovirus* in the context of its closest relatives illustrated this striking incongruence (Figure 1). In particular, both *Microhyla letovirus* and *Kanakana letovirus* were more closely related to coronaviruses in the ORF1ab region but not in the S protein where they exhibited greater sequence similarity to the *Tobaniviridae,* suggestive of past recombination events (Figure 1). In contrast, *Pacific salmon nidovirus* consistently fell as a sister-group the *Coronaviridae* in both gene regions, although with long branch lengths indicative of long periods of evolutionary divergence (Figure 1)

**Table 2.**
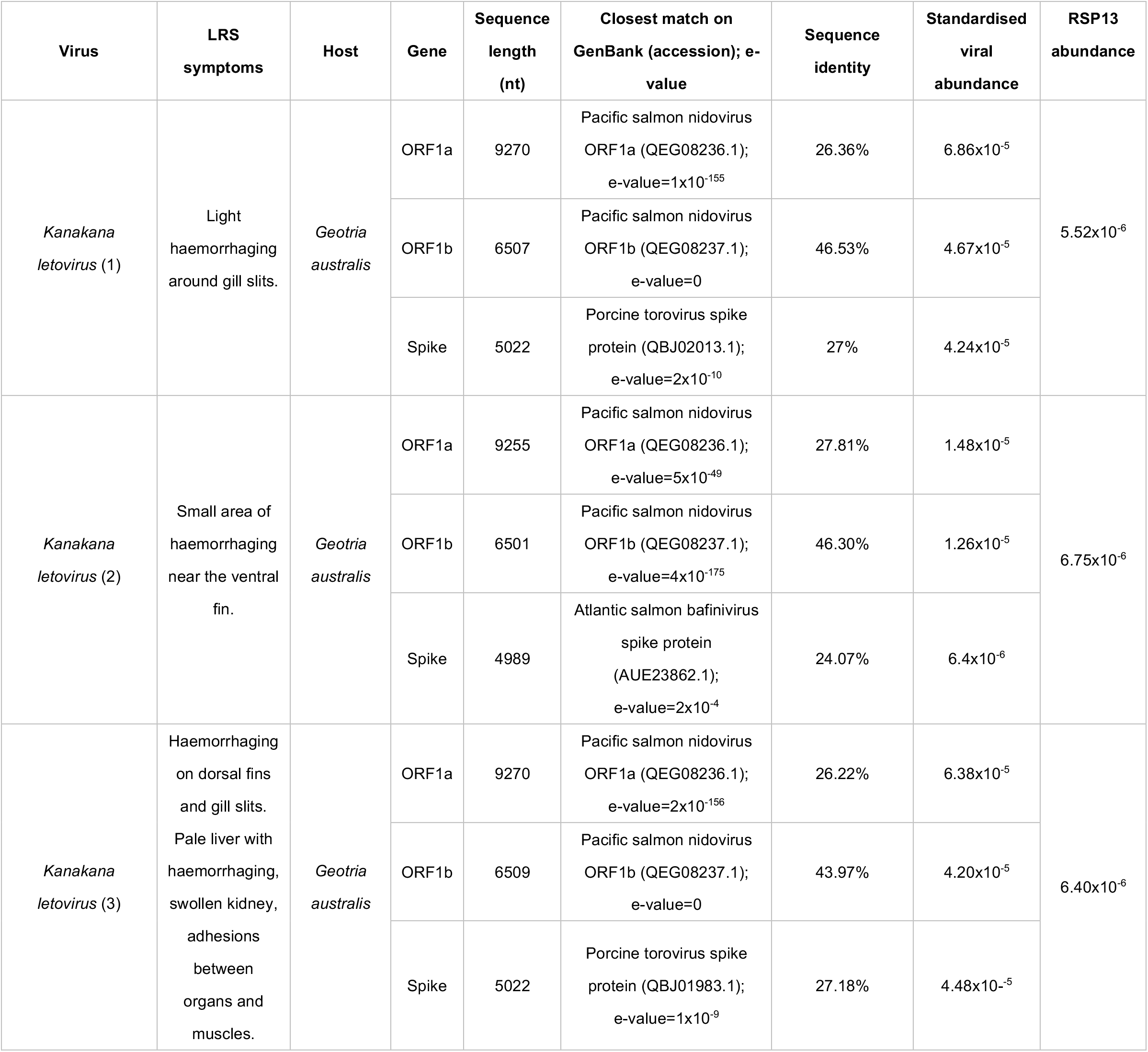
Novel virus transcripts identified in this study from pouched lamprey (*Geotria australis*), their corresponding closest genetic match on GenBank, and standardised abundances.

**Figure 1.**
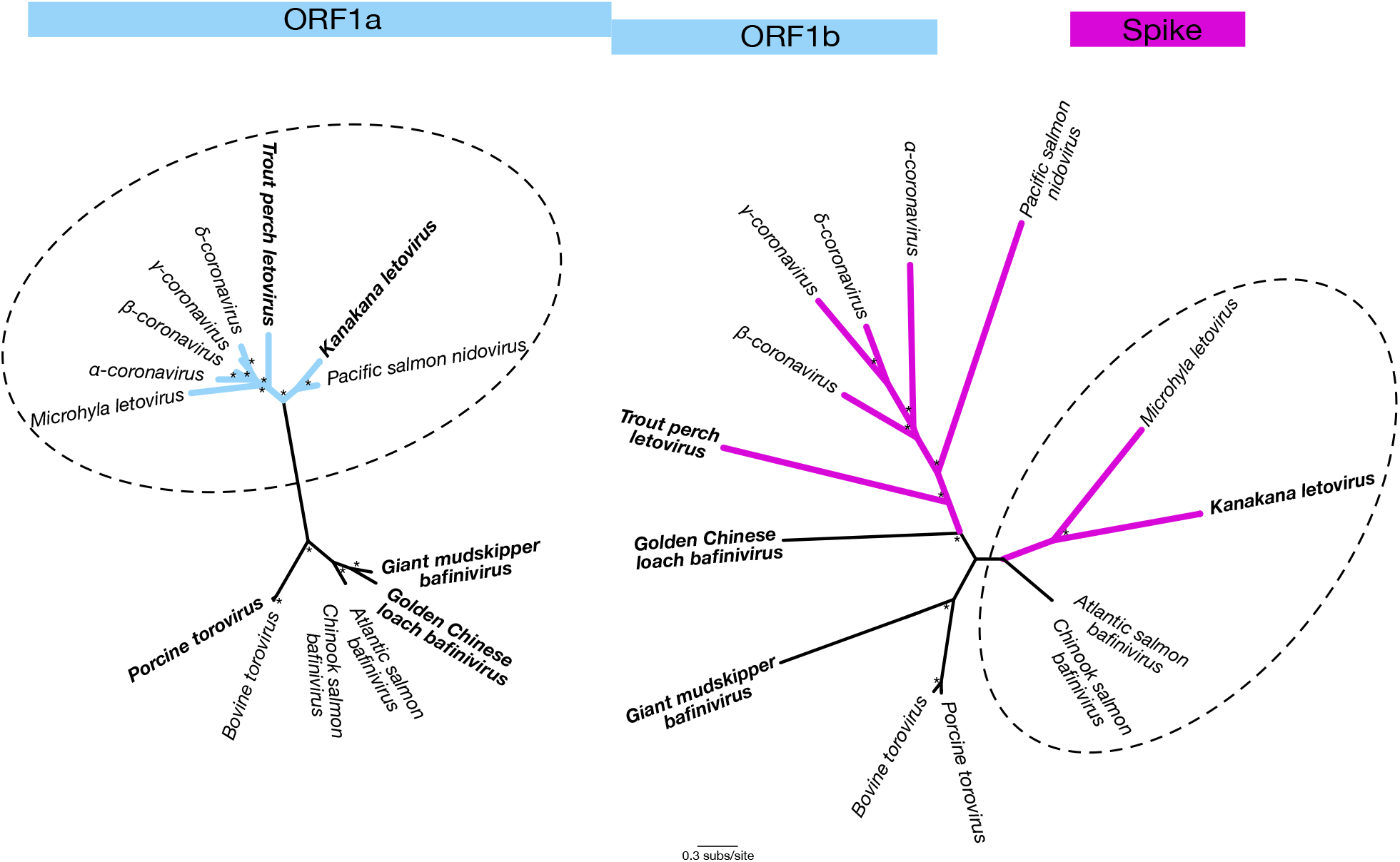
Unrooted phylogenetic trees for the open reading frame 1ab region (blue) and the spike protein (pink). Branches for viruses classified within the *Tobaniviridae* are shown in black while all other branches are coloured. A dashed line circles those taxa that are closest known relatives of *Kanakana letovirus* in each case. Four of the viruses discovered in this study in which spike proteins were identified are in bold. All branches are scaled according to the number of amino acid substitutions per site. Percentage node labels with bootstrap support >70% are indicated with an asterisk (*).

All three *Kanakana letovirus* sequencing libraries had similar sequencing depths as illustrated by the standardised abundance of the host gene, RSP13, which was detected at a lower abundance than *Kanakana letovirus* (Figure 2). However, the *Kanakana letovirus* in samples 1 and 3 were the most abundant across the ORF1ab and S proteins. The presence of *Kanakana letovirus* in seven of the individual lampreys was confirmed by RT-PCR (Supplementary Figure 1). No other viruses were detected in these data.

**Figure 2.**
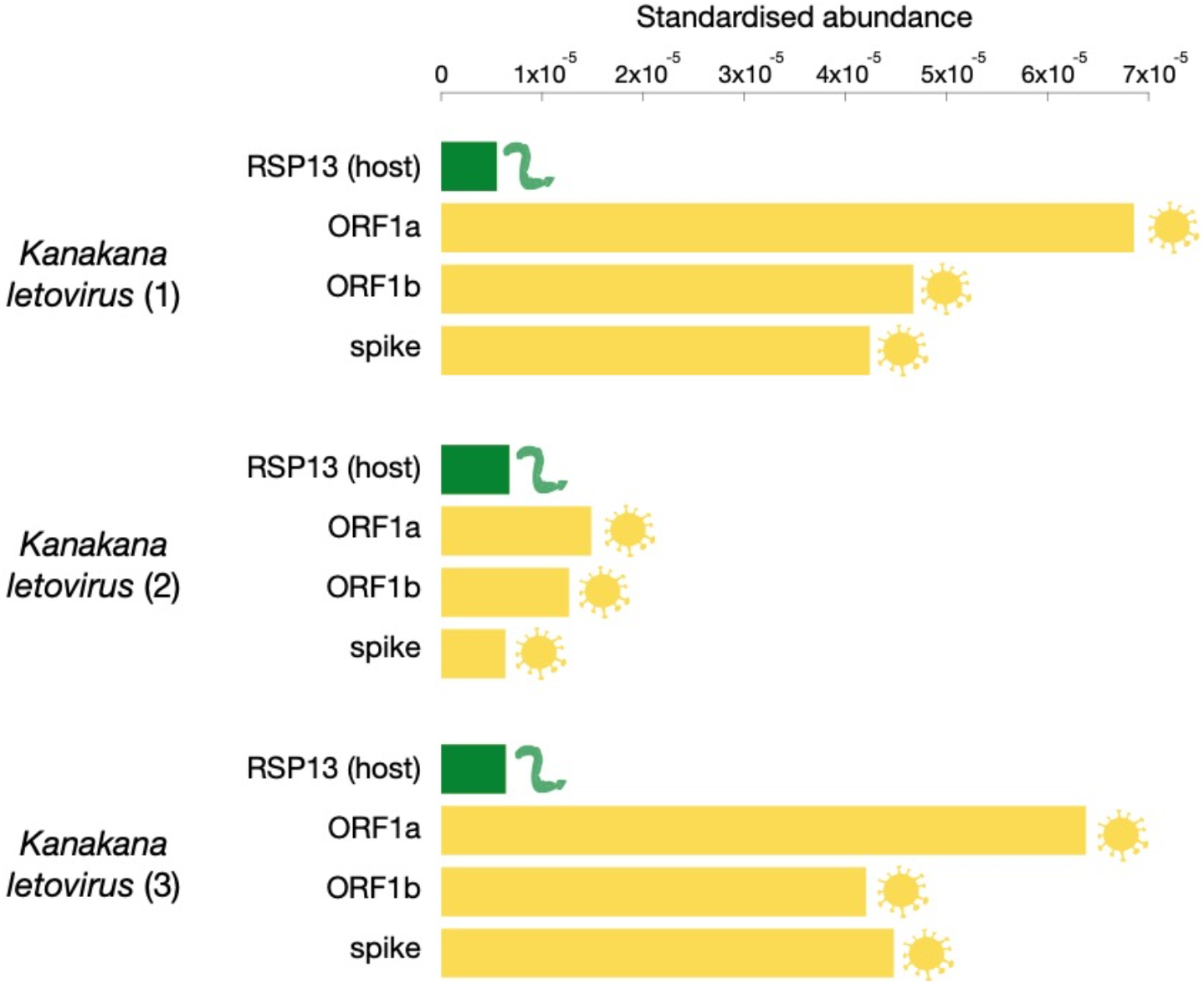
Standardised abundance of *Kanakana letovirus* transcripts (yellow) in comparison to the lamprey host gene, RSP13 (green), across the three lamprey samples that contained the highest abundance of the novel coronavirus.

### Identification of novel coronaviruses in published SRA data from bony fish

Data mining the SRA for coronavirus-like sequences revealed several potentially novel virus transcripts within sequences obtained from bony fish (Table 1). We detected coronavirus-like reads in 32 sequencing libraries (of the 1279 that met our search criteria) that were further investigated. First, we identified four new viruses that again shared OFR1b sequence similarity to *Pacific salmon nidovirus* (52-56%). These sequences were identified in blenny (*Acanthemblemaria sp*.), bluespotted mud hopper (*Boleophthalmus pectinirostris*), *Schizothorax waltoni*, which were sampled from China, as well as the broadnosed pipefish (*Syngnathus typhle*), sampled from Europe. The new viruses were tentatively named *Blenny letovirus, Mud hopper letovirus, Waltoni letovirus* and *Broadnosed pipefish letovirus*, respectively. Second, we identified a viral transcript that shared 39% amino acid sequence similarity to *Shrew coronavirus* in ORF1b and 20% amino acid sequence similarity to *Bat coronavirus* in the S protein (Table 1), although it is important to note that the vast majority of viruses in the *Letovirinae* are not yet included in the nr database. Finally, two potentially novel virus sequences were identified that, upon further inspection, shared greater sequence similarity to viruses within the *Tobaniviridae*, in a similar manner to the S protein of *Kanakana letovirus*. These two novel viruses, found in giant mudskipper (*Periophthalmodon schlosseri*) and golden Chinese loach (*Botia superciliaris*), had >50% sequence similarity in the RdRp and ~40% sequence similarity in the S protein to other fish viruses in the *Tobaniviridae* (see Table 1). These new viruses were tentatively named G*iant mudskipper bafinivirus* and *Golden Chinese loach bafinivirus*, respectively. The remaining 25 libraries that were further investigated did not possess transcripts with sequence similarity to the *Coronaviridae*.

### The evolutionary history of novel coronaviruses in aquatic vertebrates

We next inferred the evolutionary relationships of all the viruses newly identified here and related *Nidovirales* through phylogenetic analysis of amino acid sequences of ORF1b that includes the highly conserved RdRp. As noted above, two viruses identified here fell within the *Tobaniviridae*, rather than the *Coronaviridae*, and were close genetic relatives of other fish bafiniviruses (Cano et al. 2020) (Figure 3b). This analysis also revealed that fish coronaviruses fell only within the subfamily *Letovirinae* and were distinct from the *Orthocoronavirinae*. Specifically, the hosts of viruses in the *Letovirinae* comprised only aquatic vertebrates: both jawless and bony fish as well as viruses from amphibians and a loggerhead sea turtle (*Caretta caretta*) (Figure 3c).

**Figure 3.**
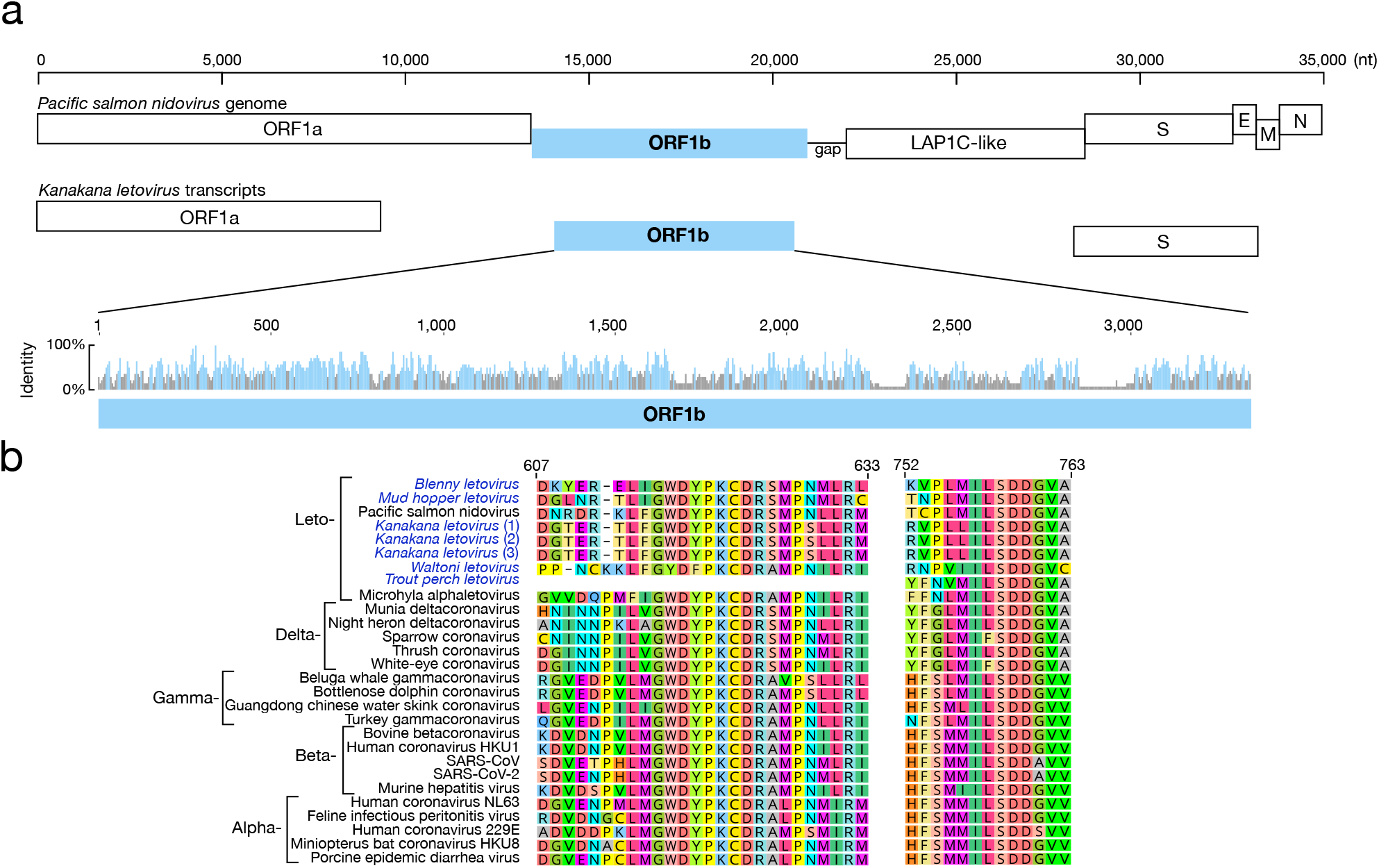
(a) Schematic phylogram showing where bony and jawless fish fall in the greater animal phylogeny and colour-coded to signify the host class in (b) and (c). (b) An order-level phylogenetic tree showing the relationships between families within the *Nidovirales*. Branches are colour-coded to signify the host class shown in part a. Branches in bold represent those viruses identified in this study. All branches are scaled according to the number of amino acid substitutions per site. (c) Maximum likelihood phylogenetic tree of the open reading frame 1b (ORF1b), which contains the viral RdRp, showing the topology of the nine newly discovered coronaviruses (bold font; blue and cyan), which fall within *Letovirinae*. The order-level tree in (b) was used to root the tree. All branches are scaled according to the number of amino acid substitutions per site. Percentage node labels with bootstrap support >70% are indicated with an asterisk (*).

Although there are broad patterns of host clustering across the coronaviruses as a whole with, for example, most of the mammalian coronaviruses falling in the genera *Alphacoronavirus* and *Betacoronavirus*, and the genus *Deltacoronavirus* comprising avian-associated viruses only, it is notable that the phylogenetic position of the *Letovirinae* does not follow that of their hosts (Figure 4a). Of particular note, *Kanakana letovirus* sampled from New Zealand pouched lamprey falls as an ingroup in this subfamily. Similar patterns can be seen for viruses in amphibians and reptiles, indicative of major cross-species transmission events. For example, the two known coronaviruses from reptiles fall across the two different sub-families, while the letoviruses identified from amphibian hosts are highly divergent, with the axolotl letovirus more closely related to viruses in fish and the frog letovirus falling in a more divergent position, forming an outgroup to other viruses in this sub-family. Indeed, phylogenetic reconciliation analysis found that host-jumping was consistently the most frequent event, although there was also clear evidence for host-virus co-divergence that likely reflects deeper branching events in the evolutionary history of these viruses (Figure 4b).

**Figure 4.**
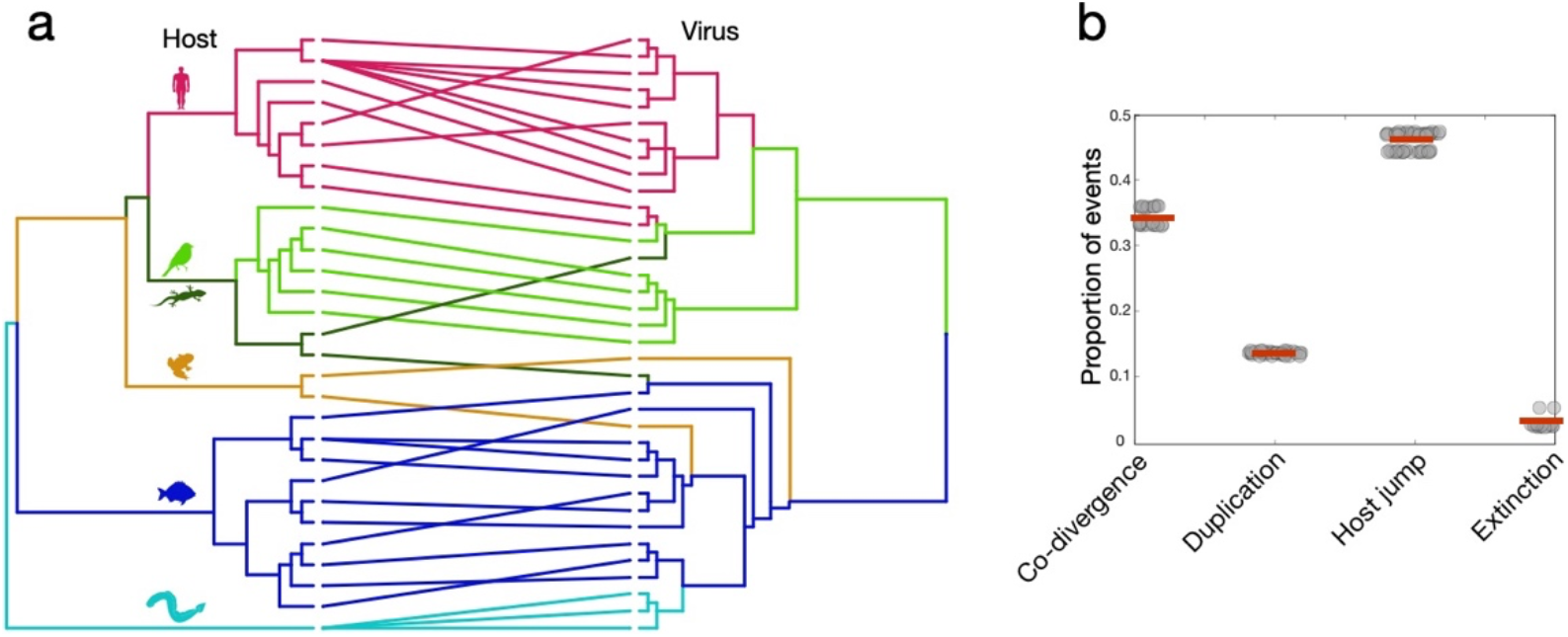
(a) Tanglegram of rooted phylogenetic trees for the *Coronaviridae* and their hosts. Lines and branches are coloured to represent host class, as in Figure 2. The ‘untangle’ function was used in TreeMap3.0 to maximise the congruence between the host (left) and virus (right) phylogenies. (b) Reconciliation analysis of each virus family using eMPRess (Santichaivekin et al. 2020) illustrates the proportion of events, with the mean indicated by a red horizontal line. The ‘event costs’ associated with incongruences between trees were conservative towards co-divergence and defined here as: 0 for co-divergence, 1 for duplication, 1 for host-jumping and 1 for extinction.

### Amino acid conservation across the Coronaviridae

Compatible with their phylogenetic relationships, the new viruses identified here possessed several motifs that are conserved among the coronaviruses, particularly within the RdRp domain in ORF1b (Figure 5). These included the amino acid sequences ‘GWDYPKCD’ and ‘ILSDDGV’ that are mostly conserved among the *Coronaviridae* but not the *Nidovirales* as a whole (Figure 5). These conserved domains are essential for metal ion chelation and binding of the primer 3′-end template complex (de Souza Luna et al. 2007).

**Figure 5.**
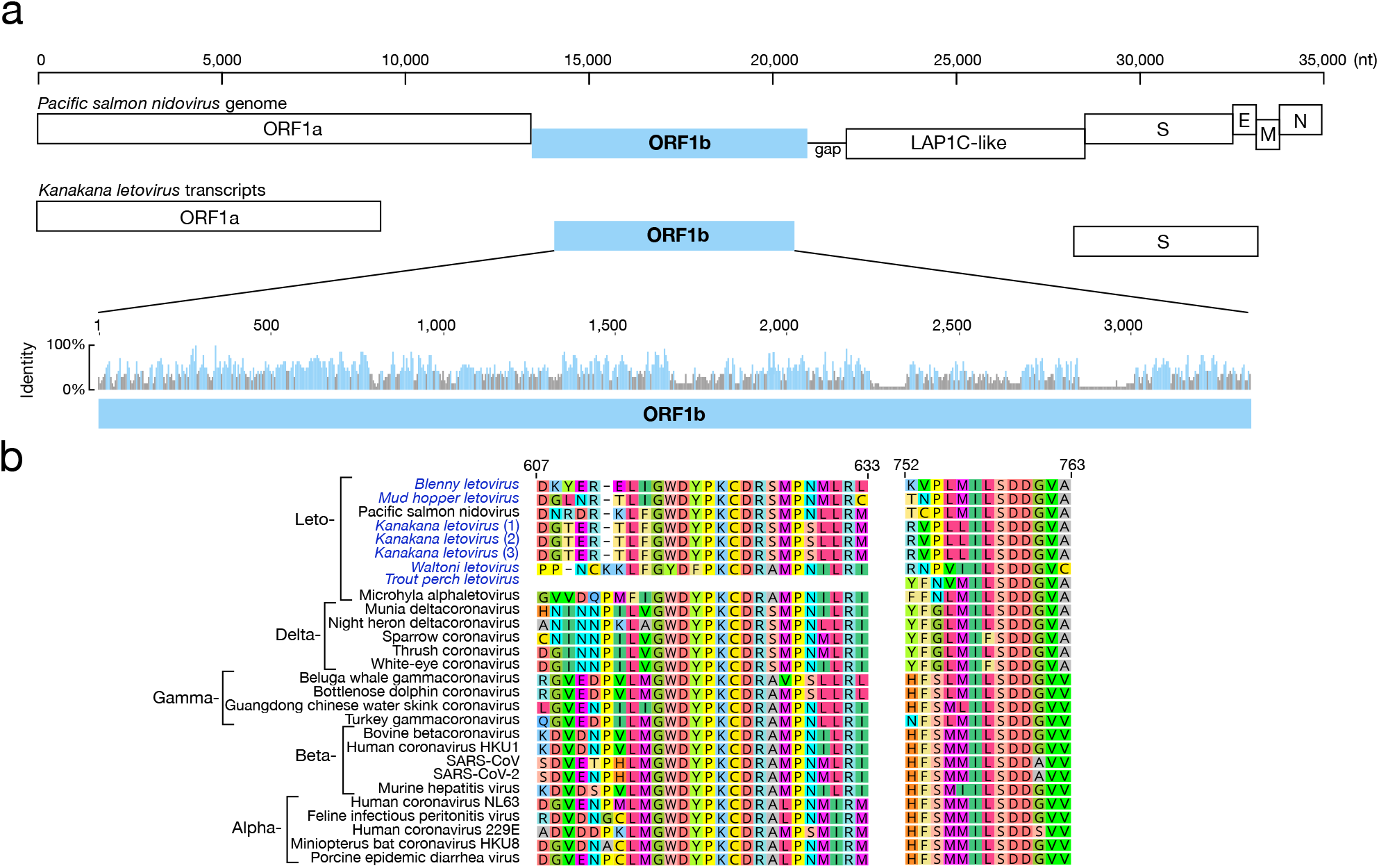
(a) Genome structure of *Pacific salmon nidovirus* and the corresponding transcripts found in the New Zealand pouched lamprey, highlighting the more conserved region, ORF1b (in blue), which includes the RdRp. A bar graph shows the sequence similarity among all viruses within the *Coronaviridae* included in Figure 3. (b) Multiple virus amino acid sequence alignment of the open reading frame 1b (ORF1b), containing the RdRp, showing motifs conserved throughout all known genera within the *Coronaviridae*, including the newly discovered viruses within the *Letovirinae* for which virus transcripts in this region were identified.

## Discussion

Virological sampling of freshwater bony and jawless fish, combined with mining publicly available sequence data, enabled us to identify nine new members of the *Nidovirales*, seven of which fell within the *Coronaviridae* and a further two fell within the related *Tobaniviridae*. The seven new coronavirus-like transcripts were close genetic relatives of viruses classified within the *Letovirinae* in the RdRp, including *Pacific salmon nidovirus*, *Microhyla letovirus* and other viruses recently identified in fishes, amphibians and reptiles, forming a sister clade to all other known coronaviruses.

The discovery of these new viruses not only expands the phylogenetic diversity of the *Letovirinae*, but it also sheds important new light on the evolution and host range in the *Coronaviridae* as a whole. In particular, our study revealed that coronavirus hosts include both freshwater and marine fish spanning multiple taxonomic orders. This includes an ancient lineage of jawless fish, herein represented by the pouched lamprey from New Zealand, that represents one of the evolutionary oldest vertebrates still to exist, and one that has remained morphologically unchanged for around 360 million years (Gess et al. 2006), although its true age has recently been debated (Miyashita et al. 2021). These jawless and bony fish hosts were also sampled across a range of geographic localities: Australia, New Zealand, China, Europe and North America. Although divergent in their sequences, conserved motifs across the entire virus family strongly supports the inclusion of these new viruses within the *Coronaviridae* and more specifically the *Letovirinae*. With the expansion of virological surveys of wildlife, it is likely that additional viruses within the *Coronaviridae* will be discovered. Indeed, the long branch lengths separating the divergent families within the *Nidovirales* is indicative of very limited sampling across this order of RNA viruses.

The genomic structure of both *Pacific salmon nidovirus* and *Ambystoma mexicanum nidovirus* includes a short, non-coding region between the ORF1ab and the remaining genes, including the S protein (Edgar et al. 2020; Mordecai et al. 2019). It has been suggested that this homologous gap represents possible genome segmentation in this clade (Edgar et al. 2020). We were, unfortunately, unable to determine the complete genomic structure of the viruses identified here and so cannot directly address this issue. It is noteworthy, however, that the S protein of *Kanakana letovirus* shared more sequence similarity with toroviruses rather than coronaviruses, while *Trout perch letovirus* and *Pacific salmon nidovirus* consistently fell within the *Coronaviridae*. This pattern of phylogenetic incongruence between these genomic regions is indicative of past recombination within the *Nidovirales*. At present, the occurrence of this process seems to be limited to only some aquatic vertebrate hosts, although given the propensity for coronaviruses to recombine (Denison et al. 2011), it is likely that this will be observed in additional species.

First recorded in 2011, lamprey populations in New Zealand’s Southland region have been affected by LRS, causing reddening along the length of the body with increased mortalities (Brosnahan et al. 2019). We cannot assign the novel *Kanakana letovirus* discovered here as the causative agent of this syndrome. Nevertheless, since lamprey are classed as a nationally vulnerable threatened species (Dunn et al. 2018), it is important that further investigation is undertaken to determine the true prevalence of coronavirus in lamprey populations and whether the virus is associated with LRS.

That *Letoviruses* form a sister clade to all other coronaviruses suggests that the backbone of the *Coronaviridae* phylogeny overall mirrors that of their vertebrate hosts, with different viral genera associated with different vertebrate classes, and the clear transition from aquatic to land, and back again (Zhang et al. 2018). Nevertheless, lamprey represent an ancient lineage of jawless fish that fall sister to all other vertebrates, including jawed fish, amphibians and reptiles. In this context it is particularly notable that the novel coronavirus identified here in the pouched lamprey does not fall basal to other fish coronaviruses, in turn revealing that cross-species transmission events have played a major role in the evolutionary history of these viruses and which is also apparent from our co-phylogenetic reconciliation analysis. Hence, as is the case with many families of RNA viruses (Geoghegan et al. 2017), the broad association between different coronaviruses and their vertebrate host classes suggests that the evolutionary history of this group likely dates to at least the origin of the vertebrates some 530 million years ago. However, on top of this overall pattern of long-term virus-host co-divergence, there have also been frequent instances of cross-species virus transmission. As the case of the pouched lamprey demonstrates, such host jumping events may be particularly commonplace in hosts that reside in aquatic environments, perhaps reflecting a combination of evolutionary antiquity (and hence more long-term opportunity for cross-species transmission) and greater physical interactions between multi-species assemblages, although this clearly needs to be investigated in greater detail.

## Materials & Methods

### Bony fish sample collection in Australia and RNA sequencing

Samples of freshwater bony fish (Osteichthyes) were collected from the Murray-Darling Basin river system in Australia between January and March 2020 and screened for coronaviruses using meta-transcriptomic sequencing (see Costa et al. (2021) for details of the broader study). Seven species of native Australian fish species were collected for this purpose: bony bream (*Nematalosa erebi*), spangled perch (*Leiopotherapon unicolor*), Australian smelt (*Retropinna semoni*), Murray-Darling rainbowfish (*Melanotaenia fluviatilis*), flat-headed gudgeon (*Philypnodon grandiceps*), western carp gudgeon (*Hypseleotris sp*.) and unspecked hardyhead (*Craterocephalus fulvus*). Three species of invasive fish were also collected: common carp (*Cyprinus carpio*), goldfish (*Carassius auratus*) and eastern mosquitofish (*Gambusia holbrooki*). Importantly, all fish species were apparently healthy, with no obvious clinical signs of disease. Fish were caught using boat electrofishing procedures and dissected immediately upon capture. Tissue specimens (liver and gills) were placed (in the same tube) in RNALater and stored in a portable −80°C freezer, then later in a −80°C freezer in the laboratory at Macquarie University, Sydney. To facilitate virus discovery, multiple individuals (1–10) were pooled according to species and the location in which they were captured. Sampling was conducted under the approval of Macquarie University Animal Ethics Committee, approval number 2019/035.

Frozen samples of liver and gill tissue were placed together in 600μl of lysis buffer containing 0.5% foaming reagent (Reagent DX, Qiagen) and 1% of ß-mercaptoethanol (Sigma-Aldrich). Submerged tissue samples were homogenized with TissueRuptor (Qiagen) for one minute at 5000 rpm. To further homogenise tissue samples and remove tissue residues, the homogenate was centrifuged at full speed for three minutes. The homogenate was carefully removed and RNA from the clear supernatant was extracted using the RNeasy Plus Mini Kit (Qiagen) following the manufacturer’s protocol. Extracted RNA was quantified using NanoDrop (ThermoFisher) and RNA from each species was pooled corresponding to the site in which they were captured, resulting in 36 sample libraries. RNA libraries were constructed using the Truseq Total RNA Library Preparation Protocol (Illumina). To enhance viral discovery and reduce the presence of non-viral reads, host ribosomal RNA (rRNA) was depleted using the Ribo-Zero-Gold Kit (Illumina) and paired-end sequencing (100bp) was performed on the NovaSeq 500 platform (Illlumina). Sample library construction, rRNA depletion and RNA sequencing were performed at the Australian Genome Research Facility (AGRF).

### Lamprey sample collection in Aotearoa New Zealand and RNA sequencing

Lamprey reddening syndrome (LRS) describes lamprey from the South Island of New Zealand that display red lesions on their gills, fins, and body walls (Brosnahan et al. 2019). From 2011 – 2013, the New Zealand Ministry of Primary Industries (MPI) received specimens exhibiting possible signs of LRS for diagnostic testing including bacteriology, histology and molecular analysis targeting specific known diseases to determine if an infectious agent was present. No infectious agent attributed to LRS was identified through this testing. Samples that MPI received in 2012 were used in the current study. These comprised 40 pouched lamprey (*Geotria australis*) (te reo Māori: kanakana - South Island, or piharau - North Island), native to New Zealand. Lamprey skin and muscle tissues displaying reddening, heart, kidney, liver, gut, and visceral fat tissues were analysed for the presence of pathogens in 2012 by MPI and these tissues were then archived in formalin-fixed paraffin-embedded (FFPE) blocks and stored at room temperature until they were sent to the University of Otago for RNA sequencing in 2020. Fresh tissues were obtained from four additional *Geotria australis* individuals, collected on separate field expeditions from the Mokau River, Aria in 2015 (n = 2) and Okuti River, Banks Peninsula in 2019 (n = 2), both of which are separate catchments areas to locations sampled in 2012. These individuals displayed no signs of LRS which has never been recorded from the Mokau or Okuti rivers. These samples were used as controls. Muscle and skin subsamples from the control individuals were taken in the field and were preserved in RNALater (Thermo Fisher Scientific) and in a −80 freezer. Sampling was conducted in consultation with local Māori (rūnanga and kaumatua).

Skin tissue was dissected from 26 FFPE blocks, representing 26 individuals. Total RNA was extracted using a Macherey-Nagel NucleoSpin^®^ totalRNA FFPE XS kit following the NucleoSpin totalRNA FFPE XS protocol. Control samples from the Mokau and Okuti rivers were extracted separately using a Zymo Research Direct-zol RNA Miniprep Plus kit. The purity of the extracted RNA from all sample types was evaluated (NanoDrop and Qubit Fluorometer) and 12 of the highest scoring samples (eight LRS and four controls) were sent to Otago Genomics Facility for sequencing using the Truseq Total RNA Library Preparation Protocol (Illumina). Host ribosomal RNA (rRNA) was depleted using the Ribo-Zero-Gold Kit (Illumina) to enhance viral discovery. Libraries were quality checked and sequenced in one lane on the HiSeq 2500 V4 Illumina platform.

### Data mining the Sequence Read Archive (SRA) for coronaviruses

To identify additional coronavirus-like sequences we screened bony fish transcriptomes present in the SRA. We focused on a subset of available SRA libraries, particularly those RNA-seq libraries without sequencing selection (e.g., poly(A)+ -selected). We excluded zebrafish (*Danio rerio*) runs from the screens due to the large number of experimental data sets. To reduce redundancy, we selected the first six runs for a given species in each experimental data set. In total, 1279 libraries (both single- and paired-end reads) were selected. SRA runs were obtained through the European Nucleotide Archive (https://www.ebi.ac.uk/ena/browser/home) that mirrors the SRA but allows for reads to be downloaded directly in a FASTQ. Reads were then analysed using our coronavirus discovery pipeline. The pipeline consisted of an initial screen that aligned the reads against a custom database of *Coronaviridae* and other protein sequences within the *Nidovirales* using Diamond BLASTx (v.2.0.2) (Buchfink et al. 2015) with an e-value threshold of 1×10^−5^. Results were manually inspected in Geneious Prime (www.geneious.com) to exclude spurious alignments to low-complexity regions. Where coronavirus-like reads were detected (n=32) we quality trimmed and assembled *de novo* raw reads into contigs using Trinity RNA-Seq (v2.9.1) (Haas et al. 2013). The assembled contigs were then compared to NCBI non-redundant protein database using Diamond BLASTx with an e-value threshold of 1×10^−5^. To identity highly divergent sequences we regularly updated our custom *Coronaviridae* protein database with the novel viruses identified. Where a coronavirus-like sequence was found we examined all SRA runs in the experimental dataset for the given species.

### Transcript sequence similarity searching for novel coronaviruses

Sequencing reads were first quality trimmed then assembled *de novo* using Trinity RNA-Seq (v.2.11.0) (Haas et al. 2013). The assembled contigs were annotated based on similarity searches against the NCBI nucleotide (nt) and non-redundant protein (nr) databases using BLASTn (Altschul et al. 1990) and Diamond BLASTx (v.2.0.2) (Buchfink et al. 2015).

### Phylogenetic analysis

To infer the evolutionary relationships of the coronaviruses, newly discovered translated viral contigs were combined with representative protein sequences from both the *Coronaviridae* and the order *Nidovirales*. All these sequences were obtained from NCBI GenBank. The sequences retrieved were then aligned with those generated here using MAFFT (v7.4) (Katoh et al. 2002) employing the E-INS-i algorithm. Ambiguously aligned regions were removed using trimAl (v.1.2) (Capella-Gutiérrez et al. 2009). To estimate phylogenetic trees, we utilised the maximum likelihood approach available in IQ-TREE (v 1.6.8) (Nguyen et al. 2015), selecting the best-fit model of amino acid substitution, LG, with ModelFinder (Kalyaanamoorthy et al. 2017), and using 1000 bootstrap replicates. Phylogenetic trees were annotated with FigTree (v.1.4.2).

To visualise the occurrence of cross-species transmission and virus-host co-divergence in the *Coronaviridae* we reconciled the co-phylogenetic relationship between coronaviruses and their hosts. A host phylogeny was manually constructed using topologies available in the appropriate literature, and a tanglegram connecting the host and virus trees was inferred using TreeMap v3.0 (Charleston 2011): lines between the trees connect the host with its virus. We utilised the ‘untangle’ function, which rotates the branches of one tree, to minimize the number of line crosses (i.e. cross-species transmission events). To quantify the relative frequencies of cross-species transmission versus virus-host co-divergence we reconciled the co-phylogenetic relationship between viruses and their hosts using eMPRess (Santichaivekin et al. 2020). This approach employs a maximum parsimony approach to determine the best ‘map’ of the virus phylogeny onto the host phylogeny. We set the duplication, host-jump and extinction event types to have a cost of one, while that of host-virus co-divergence was considered a ‘null event’ and therefore had cost of zero. Following the parsimony principle, the reconciliation proceeds by minimising the total event cost.

### Estimation of virus abundance

Transcriptomes were quantified using RNA-Seq by Expectation-Maximization (RSEM) as implemented within Trinity (Li and Dewey 2011) and standardised by the number of paired reads in a given library. This enabled us to estimate the relative abundance of each virus transcript in these data. For comparison, and to assess the sequencing depth across libraries, we also estimated the relative abundance of a stably expressed host reference gene, ribosomal protein S13 (RSP13).

### PCR confirmation

To further confirm the presence of the novel coronavirus in the pouched lamprey, total RNA was re-extracted from the 12 HiSeq-sequenced samples using the methods described above. The total RNA was then reverse transcribed (RT) using Maxima Reverse Transcriptase kit (Thermo Scientific) and ReadyMade Random Hexamers (Integrated DNA technologies [IDT]). The Maxima Reverse Transcriptase (Thermo Scientific) protocol was followed and did not include the GC-rich template options: 1 *μ*l template total RNA, 1 *μ*l ReadyMade Random Hexamers (10mM), 1 *μ*l dNTP mix (10mM), 11.5 *μ*l nuclease-free water, 4 *μ*l 5X RT Buffer, 0.5 *μ*l (20U) RiboLock RNase Inhibitor (Thermo Scientific), 1 *μ*l Maxima Reverse Transcriptase. The final volume (20 *μ*l) was stored at −20°C until used in polymerase chain reactions (PCR).

### Etymology of virus names

New viruses discovered in this study were tentatively named by drawing from their host species’ common names. The virus found in pouched lamprey used the Māori name, kanakana, since they were identified in the South Island of New Zealand and following discussions with representatives for iwi (Māori tribes). We suggest that all of these viruses be grouped within the subfamily *Letovirinae*.

## Data availability

Sequencing reads and virus sequences from pouched lamprey samples are available online via the Genomics Aotearoa database to maintain Māori data sovereignty of native species (taonga). All other data are available on NCBI (accession numbers available soon).

## Supplementary Material

**Supplementary Figure 1**. Agarose gels electrophoresis showing PCR products from primers that targeted regions in coronavirus ORF1ab (including the RdRp) for 12 individuals of pouched lamprey.

**Supplementary Table 1**. Primer set used for the RT-PCR confirmation of *Kanakana letovirus* in specimens of pouched lamprey.

## Acknowledgements

The authors thank Peter Stockman (Ngati Maniapoto, Te Rūnanga o Wairewa), and Keith Bradshaw and Duncan Ryan (Te Runaka o Awarua) for sample collection. We are grateful to the iwi representatives for providing insights and for their collaboration, and to the New Zealand Ministry for Primary Industries for access to samples.

