## Supplementary Figure 1 for "Slippery when wet: cross-species transmission of divergent coronaviruses in bony and jawless fish and the evolutionary history of the *Coronaviridae*"

**Supplementary Figure 1**. Agarose gels electrophoresis showing PCR products from three sets of primers that targeted regions in coronavirus ORF1b (including the RdRp) for pouched lamprey.

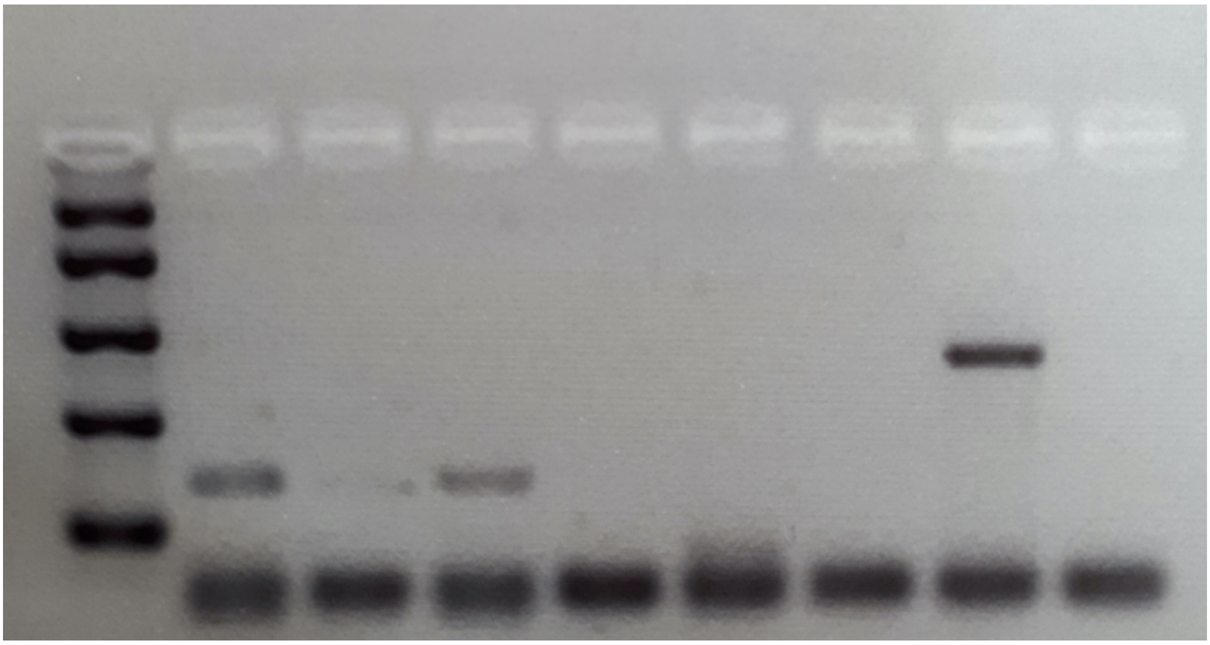
