## Supplementary Table 1 for "Slippery when wet: cross-species transmission of divergent coronaviruses in bony and jawless fish and the evolutionary history of the *Coronaviridae*"

**Supplementary Table 1**. List of primer sets used for the RT-PCR confirmation of *Kanakana letovirus* in specimens of pouched lamprey.

| Transcript region (bp) | Product (bp) | 5' Forward 3' | 5' Reverse 3' | Tm (C) |
| --- | --- | --- | --- | --- |
| 1179 to 1309 | 150 | TCATGGCCGTGAACAAACGA | CGAAGAAGGCTCGGCATTGA | 60 |
